# An analog to digital converter controls bistable transfer competence development of a widespread bacterial integrative and conjugative element

**DOI:** 10.1101/2020.05.27.119875

**Authors:** Nicolas Carraro, Xavier Richard, Sandra Sulser, François Delavat, Christian Mazza, Jan Roelof van der Meer

## Abstract

Conjugative transfer of the integrative and conjugative element ICE*clc* in *Pseudomonas* requires development of a transfer competence state in stationary phase, which arises only in 3-5% of individual cells. The mechanisms controlling this bistable switch between non-active and transfer competent cells have long remained enigmatic. Using a variety of genetic tools and epistasis experiments in *P. putida*, we uncovered an ‘upstream’ cascade of three consecutive transcription factor-nodes, which controls transfer competence initiation. Initiation activates a feedback loop, which stochastic modeling and experimental data demonstrated acts as a scalable converter of unimodal input to bistable output. The feedback loop further enables prolonged production of a transcription factor that ensures ‘downstream’ transfer competence formation in activated cells. Phylogenetic analyses showed that the ICE*clc* regulatory factors are widespread among *Gamma-* and *Beta*-proteobacteria, highlighting its evolutionary conservation and prime importance to control the behaviour of this wide family of conjugative elements.

## Introduction

Biological bistability refers to the existence of two mutually exclusive stable states within a population of genetically identical individuals, leading to two distinct phenotypes or developmental programs^1^. The basis for bistability lies in a stochastic regulatory decision resulting in cells following one of two possible specific genetic programs that determine their phenotypic differentiation^2^. Bistability has been considered as a bet-hedging strategy leading to an increased fitness of the genotype by ensuring survival of one of both phenotypes depending on environmental conditions^3^. A number of bistable differentiation programs is well known in microbiology, notably competence formation and sporulation in *Bacillus subtilis*^4,5^, colicin production and persistence in *Escherichia coli*^6^, virulence development of *Acinetobacter baumannii*^7^, or the lysogenic/lytic switch of phage lambda^8,9^.

The dual lifestyle of the *Pseudomonas* integrative and conjugative element (ICE) ICE*clc* has also been described as a bistable phenotype (Fig. 1A)^10^. In the majority of cells ICE*clc* is maintained in the integrated state, but a small proportion of cells (3-5%) in stationary phase activates the ICE transfer competence program^10,11^. Upon resuming growth, transfer competent (tc) donor cells excise and replicate the ICE^12^, which can conjugate to a recipient cell, where the ICE can integrate^11^. ICE*clc* transfer competence comprises a differentiated stable state, because initiated tc cells do not transform back to the ICE-quiescent state. Although tc cells divide a few times, their division is compromised by the ICE and eventually arrests completely^13,14^.

**Fig 1.**
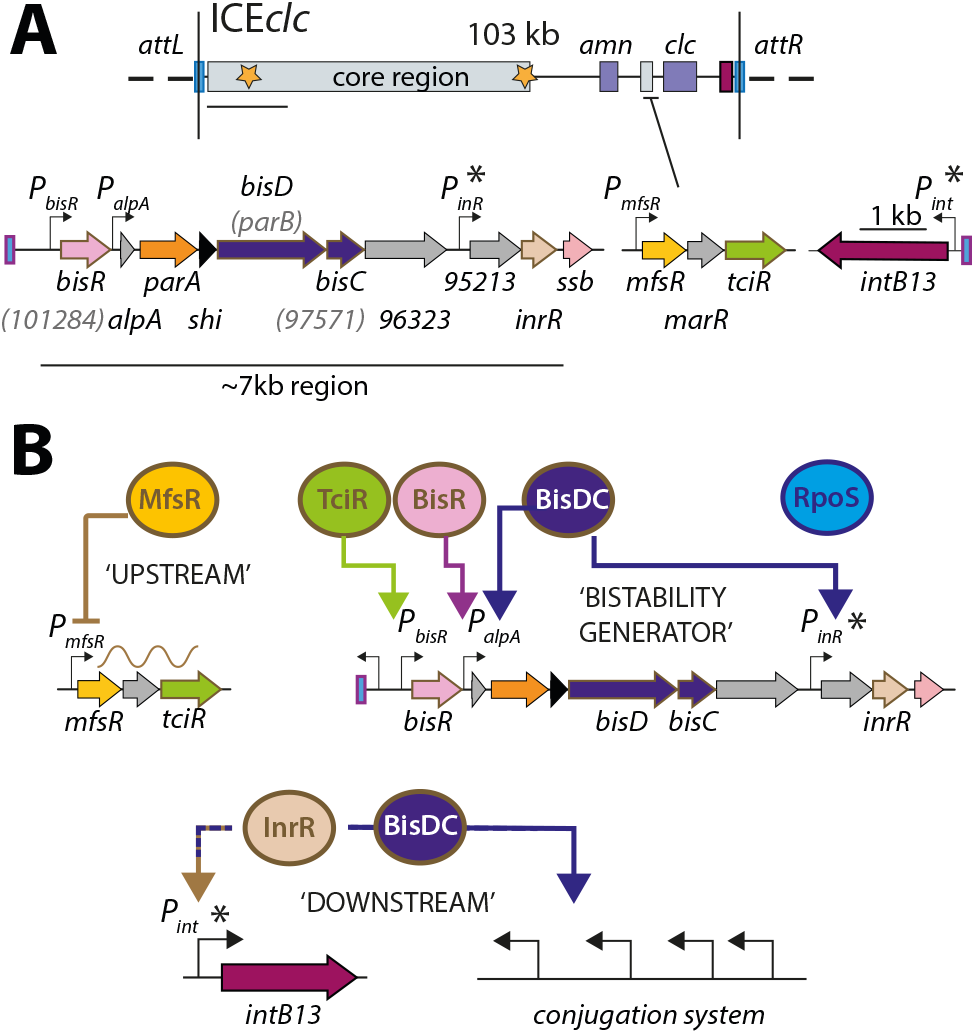
ICE*clc* and postulated regulation network for transfer competence formation. **A.** Schematic representation of the genetic organization of ICE*clc* (GenBank accession number AJ617740.2). Loci of interest are detailed below the general map and drawn to scale. Note the ~7-kb left-end region, which is the major focus of the study. Genes are represented by colored arrows with their name below (former names shown in lighter font inside brackets). Promoters are represented by hooked arrows pointing towards the transcription orientation. Those marked with an asterisk are expressed only in the subpopulation of transfer competent cells. *attL* and *attR,* attachment sites; *clc* genes: 3-chlorocatechol degradation, *amn* genes: 2-aminophenol degradation. **B.** Postulated model of ICE*clc* transfer competence regulation. An ‘upstream’ part of MfsR autorepressing its own transcription and that of *tciR*^26^; TciR activating *bisR* expression and BisR activating the *alpA* promoter. The ‘bistability generator’ of the feedback loop involving BisDC and *P_alpA_*. The ‘downstream’ parts activated by BisDC and implying additional factors RpoS and InrR^10,33^ in the subpopulation of transfer competent cells (*P_inR_, P_int_*).

ICEs have attracted wide general interest because of the large variety of adaptive functions they can confer to their host, including resistance to multiple antibiotics^15–17^, or metabolism of xenobiotic compounds, such as encoded by ICE*clc*^18,19^. Although the existence and the fitness consequences of the ICE*clc* bistable transfer competence pathway have been studied in quite some detail, the regulatory basis for its activation has remained largely elusive^20^. In terms of its genetic makeup, ICE*clc* seems very distinct from the well-known SXT/R391 family of ICEs^21^ and from ICE*St1*/ICE*St3* of *Streptococcus thermophilus*^22^. These carry analogous genetic regulatory circuitry to the lambda prophage, which is characterized by a typical double-negative feedback control^23,24^. Transcriptomic studies indicated that about half of the ICE*clc* coding capacity (~50 kb) is higher expressed in stationary than exponential phase cultures grown on 3-chlorobenzoate (3-CBA), and organized in at least half a dozen transcriptional units^25^. A group of three consecutive regulatory genes precludes ICE*clc* activation in exponentially growing cells, with the first gene (*mfsR*) constituting a negative autoregulatory feedback (Fig. 1B)^26^. Overexpression of the most distal of the three genes (*tciR*), leads to a dramatic increase of the proportion of cells activating the ICE*clc* transfer competence program^26^. Despite this initial discovery, however, the nature of the regulatory network architecture leading to bistability and controlling further expression of the ICE*clc* genes in tc cells has remained enigmatic.

The primary goal of this work was to dissect the regulatory factors and nodes underlying the activation of ICE*clc* transfer competence. Secondly, given that transfer competence only arises in a small proportion of cells in a population, we aimed to understand how the regulatory architecture yields and maintains ICE bistability. We essentially followed two experimental strategies and phenotypic readouts. First, known and suspected regulatory elements were seamlessly deleted from ICE*clc* in *P. putida* and complemented with inducible plasmid-cloned copies to study their epistasis in transfer of the ICE. Secondly, individual and combinations of suspected regulatory elements were expressed in a *P. putida* host without ICE, to study their capacity to activate the ICE*clc* promoters *P_int_* and *P_inR_*, which normally only express in wild-type tc cells (Fig. 1)^10^. As readout for their activation we quantified fluorescent reporter expression from single copy chromosomally integrated transcriptional fusions, as well as the proportion of cells expressing the reporters using quantile-quantile plotting as previously described^27^. On the basis of the discovered key regulators and nodes, we then developed a conceptual mathematical model to show by stochastic simulations how bistability is generated and maintained. This suggested that the ICE*clc* transfer competence regulatory network essentially converts an analog input signal from the ‘upstream’ regulatory branch (Fig. 1B) to a digital (bistable) output, in a scalable manner. We experimentally verified this scalable analog-digital conversion in a *P. putida* without ICE*clc* but with the reconstructed bistability generator. The key ICE*clc* bistability regulatory elements are conserved among a wide range of Proteobacteria, illustrating their importance for the behavior of this conglomerate of related ICEs.

## Results

### Activation of ICE*clc* starts with the LysR-type transcription regulator TciR

Previous work had implied an ICE*clc*-located operon of three consecutive regulatory genes (*mfsR, marR* and *tciR*, Fig. 1) in control of transfer competence formation^26^. That work had shown that *mfsR* codes for an autorepressor, whose deletion yielded unhindered production of the LysR-type activator TciR. As a result, the proportion of tc cells is largely increased in *P. putida* UWC1 bearing ICE*clc-ΔmfsR*^11, 26^. We reproduced this state of affairs here by cloning *tciR* under control of the IPTG-inducible *P_tac_* promoter on a plasmid (pME*tciR*) in *P. putida* UWC1-ICE*clc*. In absence of cloned *tciR*, transfer of wild-type ICE*clc* from succinate-grown *P. putida* to an ICE*clc*-free isogenic *P. putida* was below detection limit, indicating that spontaneous ICE activation under those conditions is negligible (Fig. 2A). In contrast, inducing *tciR* expression by IPTG addition triggered ICE*clc* transfer from succinate-grown cells up to frequencies close to those observed under wild-type growth conditions with 3-CBA^28^ (10^−2^ transconjugant colony-forming units (CFU) per donor CFU, Fig. 2A). Transfer frequencies were lower in the absence of IPTG indicating that leaky expression of *tciR* from *P_tac_* was sufficient to trigger ICE*clc* transfer (Fig. 2A). These results confirmed the implication of TciR and thus we set out to identify its potential activation targets on ICE*clc*.

**Fig 2.**
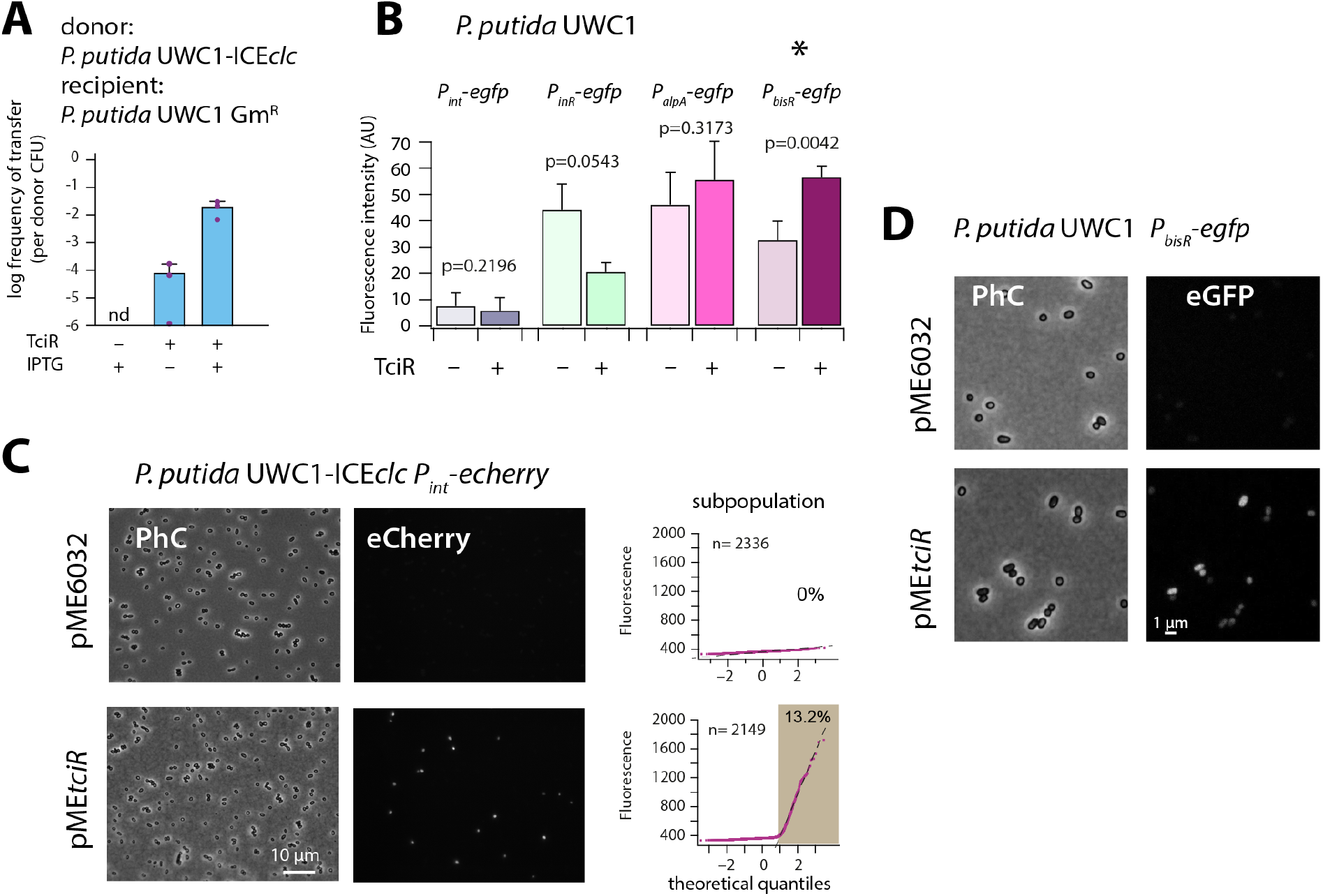
The LysR-type regulator TciR links to a single node in the regulation network. **A.** Ectopic overexpression of *tciR* induces ICE*clc* wild-type conjugative transfer under non-permissive conditions. Bars show the means (+ one standard deviation) of transconjugant formation after 48 h in triplicate matings using *P. putida* UWC1 donors carrying the indicated ICE*clc* or plasmids, in absence (-) or presence (+) of 0.1 mM IPTG, and with a Gm^R^-derivative of *P. putida* as recipient. Dots represent individual transfer; nd: not detected (<10^−7^ for the three replicates). **B.** Reporter expression from single copy chromomosomal *P_bisR_*, *P_inR_*, *P_int_*, or *P_alpA_* transcriptional *egfp* fusions in *P. putida* UWC1 without ICE*clc* as a function of ectopically expressed TciR, in comparison to strains carrying the empty vector pME6032. Bars show means of the 75^th^ percentile fluorescence of 500–1000 individual cells each per triplicate culture grown on succinate, induced with 0.05 mM IPTG. Error bars denote standard deviation from the means from biological triplicates. AU, arbitrary units of brightness at 500 ms exposure. p-values derive from pair-wise comparisons in t-tests between cultures expressing TciR and not. **C.** Proportion of cells expressing eCherry from a single-copy chromosomal insertion of *P_int_* in *P. putida* with ICE*clc* in presence of induced TciR (pME*tciR*, 0.05 mM IPTG) or with empty vector (pME6032). Fluorescence images scaled to same brightness (300–2000). Diagrams show quantile-quantile plots of individual cell fluorescence levels, with *n* denoting the number of analyzed cells and the shaded part indicating the subpopulation size expressing *P_int_-echerry.* **D.** Fluorescence images of *P. putida* without ICE*clc* with a single-copy chromosomal *P_bisR_*-*egfp* fusion in presence of empty vector or of induced TciR. Images scaled to same brightness (300–1200).

Induction of *tciR* from pME*tciR* in *P. putida* without ICE*clc* was insufficient to trigger eGFP production from a single-copy *P_int_* promoter, which is a hallmark of induction of ICE*clc* transfer competence (Fig. 2B)^10,11^. In contrast, in presence of ICE*clc*, similar induction of *tciR* yielded a clear increased subpopulation of activated cells (Fig. 2C). This suggested, therefore, that TciR does not directly activate *P_int_*, but only through one or more other ICE-located factors. To search for such potential factors, we examined in more detail the genes in a 7-kb region at the left end of ICE*clc* (close to the *attL* site, Fig. 1A), where transposon mutations had previously been shown to influence *P_int_* expression^29^. In addition, three promoters had been characterized in this region (Fig. 1A)^25^, which we tested individually for potential activation by TciR (Fig. 2B).

Promoters were fused with a promoterless *egfp* gene and inserted in single copy into the genome of *P. putida* UWC1 without ICE*clc* (Materials and Methods). Induction of *tciR* from *P_tac_* on pME*tciR* did not yield any eGFP fluorescence in *P. putida* UWC1 containing a single-copy *P_alpA_*- or *P_inR_*-*egfp* transcriptional fusion (Fig. 2B). In contrast, the *P_bisR_-egfp* fusion was activated upon induction of TciR compared to a vector-only control (p = 0.0042, paired t-test, Fig. 2B, D). This suggested that the link between TciR and ICE*clc* transfer competence proceeds through transcription activation of the promoter upstream of the gene *bisR* (previously designated *orf101284*). This transcript has previously been mapped and covers a single gene^25^. We renamed this gene as *bisR*, or bistability regulator, for its presumed implication in ICE*clc* <u>bis</u>tability control (Fig. 1B, see further below).

### BisR is the second step in the cascade of ICE*clc* transfer competence initiation

*bisR* is predicted to encode a 251-aa protein of unknown function with no detectable Pfam-domains. Further structural analysis using Phyre2^30^ suggested three putative domains with low confidence (between 38% and 53%, Fig. S1). One of these is a predicted DNA-binding domain, which hinted at the possible function of BisR as a transcriptional regulator itself. BlastP analysis showed that BisR homologs are widely distributed and well conserved among *Beta-, Alpha-* and *Gammaproteobacteria*, with homologies ranging from 43–100% amino acid identity over the (quasi) full sequence length (Fig. S2).

In order to investigate its potential regulatory function, *bisR* was cloned on a plasmid (pME*bisR*) and introduced into *P. putida* UWC1-ICE*clc*. Inducing *bisR* by IPTG addition from *P_tac_* triggered high rates of ICE*clc* transfer on succinate media (Fig. 3A). Deletion of *bisR* on ICE*clc* abolished its transfer, even upon overexpression of *tciR*, but could be restored upon ectopic expression of *bisR* (Fig. 3A). This showed that the absence of transfer was due to the lack of intact *bisR*, and not to a polar effect of *bisR* deletion on a downstream gene (Fig. 1A). In addition, transfer of an ICE*clc* deleted for *tciR*^26^ could be restored by ectopic *bisR* expression (Fig. 3A). This indicated that TciR is ‘upstream’ in the regulatory cascade of BisR, and that TciR does not act anywhere else on the expression of components crucial for ICE*clc* transfer.

**Fig 3.**
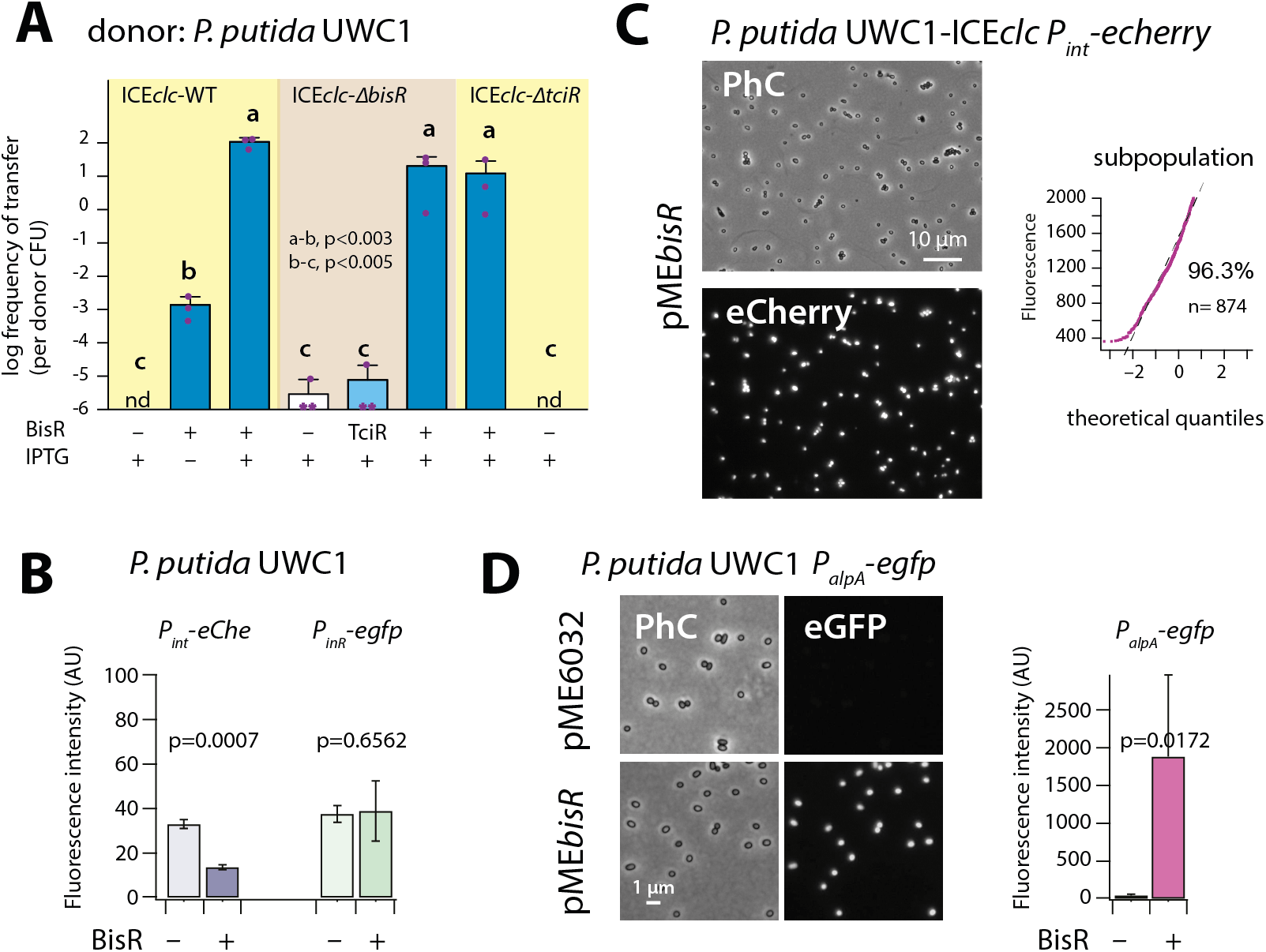
Identification of BisR as a new intermediary regulator for *P_alpA_* activation. **A.** Ectopic overexpression of *bisR* induces ICE*clc* conjugative transfer under non-permissive conditions and from ICE*clc* deleted of key regulatory genes. For explanation of bar diagram meaning, see Fig. 2A legend. BisR (+), plasmid with *bisR*; TciR, pME*tciR*; –, empty vector pME6032. nd: not detected (<10^−7^ for the three replicates). Letters indicate significance groups in ANOVA followed by post-hoc Tukey testing. **B.** Absence of direct induction by BisR of *P_int_* or *P_inR_* fluorescence reporters in *P. putida* without ICE. For explanation of bars, see Fig. 2B legend. **C.** Population-wide expression of *P_int_-echerry* in *P. putida* with ICE*clc* upon ectopic induction of plasmid-located BisR (pME*bisR*, 0.05 mM IPTG). Image brightness scale: 300– 2000. For vector control, see Fig. 2C. **D.** Induced BisR from plasmid leads to reporter expression from the *alpA*-promoter in all cells of *P. putida* without ICE*clc*. Image brightness scales: 300–1200. Bars show means and standard deviation from median fluorescence intensity of single cells (*n* = 500–1000, summed from 6-12 images per replicate) of biological triplicates. p-value derives from pair-wise t-test between cultures with empty vector (–) and those with induced BisR (+).

IPTG induction of *bisR* in *P. putida* without ICE again did not yield activation of the single-copy *P_int_* or *P_inR_* transcriptional reporter fusions (Fig. 3B). In contrast, BisR induction in *P. putida* UWC1 with ICE*clc* led to a massive activation of the same reporter constructs in virtually all cells (Fig. 3C), compared to a vector-only control (Fig. 2C, pME6032). This suggested that BisR was an(other) intermediate regulator step in the complete cascade of activation of ICE*clc* transfer competence. Of the tested ICE–promoters within this 7-kb region, BisR induction triggered very strong expression from a single copy *P_alpA_*–*egfp* transcriptional fusion in all cells (Fig. 3D). This indicated that BisR is a transcription activator, and an intermediate regulator between TciR and further factors encoded downstream of the *alpA-* promoter (Fig. 1B).

### A new regulator BisDC is the last step in the activation cascade

Next, we thus focused our attention on the genes downstream of the *alpA-*promoter. Cloning the genes from *alpA* all the way to *inrR* (Fig. 1A) on plasmid pME6032 under control of *P_tac_* and inducing that construct with IPTG resulted in activation of *P_inR_–egfp* and *P_int_*–*echerry* expression in *P. putida* without ICE*clc* (Fig. 4A). Both these promoters had been silent upon activation of TciR or BisR (Fig. 2B, 3B). This indicated that one or more regulatory factors directly controlling expression of *P_inR_* and/or *P_int_* were encoded in this region, which we tried to identify by subcloning different gene configurations.

**Fig 4.**
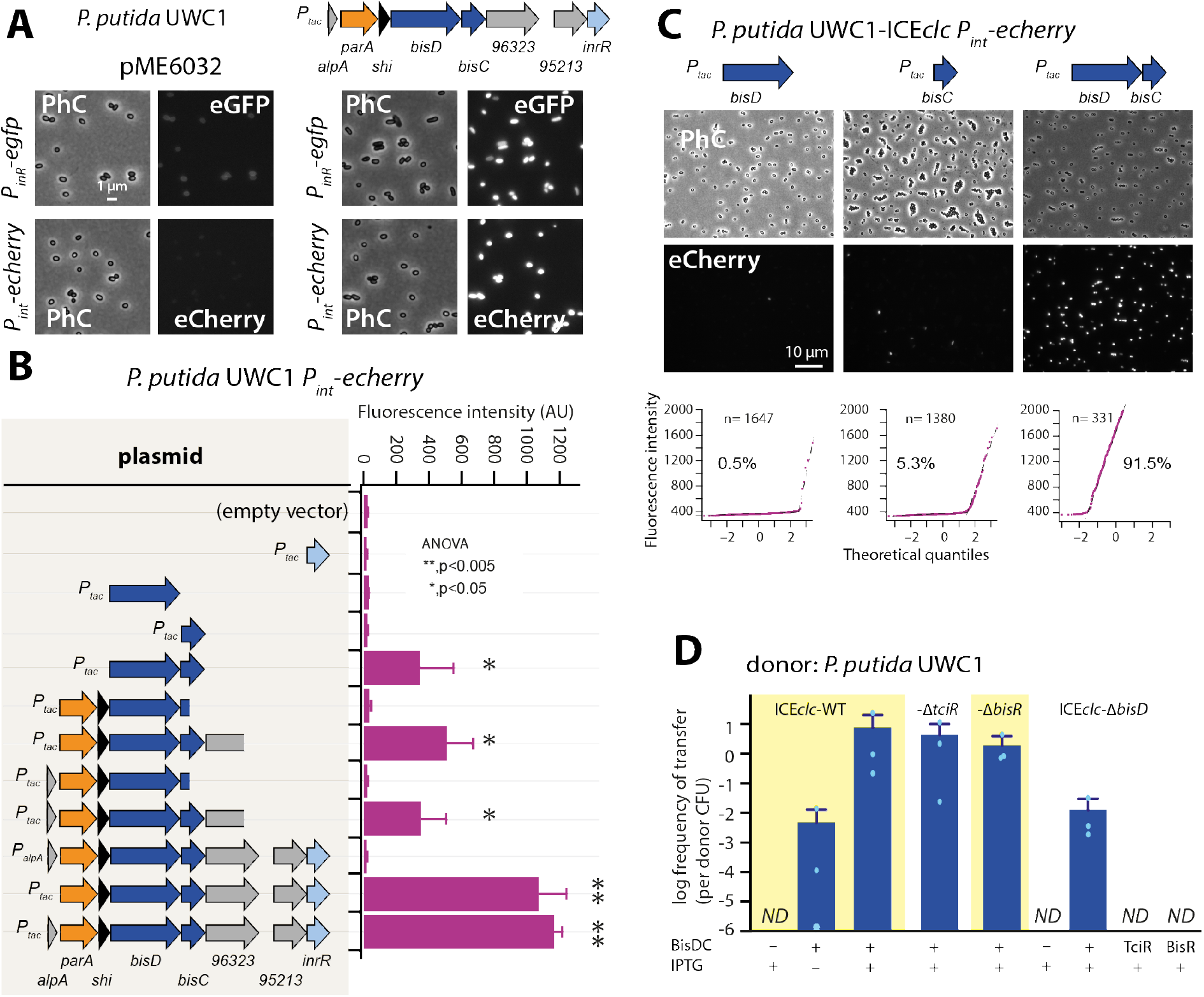
A new regulatory factor BisDC for activation of downstream ICE*clc* functions. **A.** IPTG (0.05 mM) induction of a plasmid with the cloned ICE*clc* left-end gene region (as depicted on top) leads to reporter expression from the ‘downstream’ *P_inR_-* and *P_int_*-promoters in *P. putida* without ICE*clc*. Fluorescence images scaled to same brightness (300–1200). **B.***P_int_*-*echerry* reporter expression upon IPTG induction (0.05 mM) of different plasmid-subcloned left-end region fragments (grey shaded area on the left) in *P. putida* without ICE*clc.* Bars show means of median cell fluorescence levels with one standard deviation, from triplicate biological cultures (*n* = 500–1000 cells, summed from 6–12 images per replicate). Asterisks denote significance groups in ANOVA followed by post-hoc Tukey testing. **C.** Population response of *P_int_*-*echerry* induction in *P. putida* with ICE*clc* in presence of plasmid constructs expressing *bisD*, *bisC* or both (fluorescence images scaled to 300–2000 brightness). Quantile-quantile plots (n = number of cells) below show the estimated size of the responding subpopulation. **D.** Effect of *bisDC* induction from cloned plasmid (0.05 mM IPTG) on conjugative transfer of ICE*clc* wild-type or mutant derivatives. Transfer assays as in legend to Fig. 2A. *ND*, below detection limit (10^−7^).

Removing *alpA* from the initial construct had no measurable effect on expression of the fluorescent reporters, but replacing *P_tac_* by the native *P_alpA_* promoter abolished all *P_int_* reporter activation (Fig. 4B). This suggested that *P_alpA_* is silent without activation by BisR (see below) and no spontaneous production of regulatory factors occurred. Removing three genes at the 3’ extremity (i.e., *orf96323*, *orf95213* and *inrR*) reduced *P_int_–echerry* reporter expression, but a fragment with a further deletion into the *bisC* gene was unable to activate *P_int_* (Fig. 4B). Induction of *inrR* alone did not result in *P_int_* activation (Fig. 4B). Deletion of *parA* and *shi* at the 5’ end of the fragment still enabled reporter expression from *P_int_*, narrowing the activator factor regions down to two genes, previously named *parB* and *orf97571,* but renamed here to *bisD* and *bisC* (Fig. 4B). Neither *bisC* or *bisD* alone, but only the combination of *bisDC* resulted in reporter expression from *P_int_* in *P. putida* UWC1 without ICE*clc* (Fig. 4B), and similarly, of *P_inR_* (Fig. S3). In the presence of ICE*clc*, inducing either *bisC* or *bisD* from a plasmid yielded a small proportion of cells expressing the *P_int_* reporters (Fig. 4C), which was absent in a *P. putida* carrying an ICE*clc* with a deletion of *bisD* (Fig. S4). In contrast, induction of *bisDC* in combination caused a majority of cells to express fluorescence from *P_int_* in *P. putida* containing ICE*clc* (Fig. 4C) or ICE*clc-ΔbisD* (Fig. S4). These results indicated that BisDC acts as an ensemble to activate transcription, and this pointed to *bisDC* as the last step in the regulatory cascade, since it was the minimum unit sufficient for activation of the *P_int_–*promoter, which is exclusively expressed in the subpopulation of tc cells of wild-type *P. putida* with ICE*clc*^11^.

Induction of *bisDC* from plasmid pME*bisDC* yielded high frequencies of ICE*clc* transfer from *P. putida* UWC1 under succinate-growth conditions (Fig. 4D). Expression of BisDC also induced transfer of ICE*clc-*variants deleted for *tciR* or for *bisR* (Fig. 4D). This confirmed that both *tciR* and *bisR* relay activation steps to *P_bisR_* and *P_alpA_*, respectively, but not to further downstream ICE promoters (Fig. 1B). Moreover, an ICE*clc* deleted for *bisD* could not be restored for transfer by overexpression of *tciR* or *bisR*, but only by complementation with *bisDC* (Fig. 4D). Interestingly, the frequency of transfer of an ICE*clc* lacking *bisD* complemented by expression of *bisDC* in *trans* was two orders of magnitude lower than that of similarly complemented wild-type ICE*clc*, ICE*clc* with *tciR-* or *bisR-*deletion (Fig. 4D). This was similar as the reduction in reporter expression observed in *P. putida* ICE*clc-ΔbisD* complemented with pME*bisDC* compared to wild-type ICE*clc* (Fig. S4), and suggested the necessity of some ‘reinforcement’ occurring in the wild-type configuration that was lacking in the *bisD* deletion and could not be restored by *in trans* induction of plasmid-cloned *bisDC*.

### BisDC is part of a positive autoregulatory feedback loop

To investigate this potential ‘reinforcement’ in wild-type configuration, we revisited the potential for activation of the *alpA* promoter. Induction by IPTG of the plasmid-cloned fragment encompassing the gene region *parA-shi-bisDC* caused strong activation of reporter gene expression from *P_alpA_* in *P. putida* without ICE*clc* (Fig. 5A). The minimal region that still maintained *P_alpA_* induction encompassed *bisDC*, although much lower than with a cloned *parA-shi-bisDC* fragment (Fig. 5A). Interestingly, when the *parA-shi-bisDC* fragment was extended by *alpA* itself, reporter expression from *P_alpA_* was abolished, whereas also a fragment containing only *alpA* caused significant repression of the *alpA* promoter (Fig. 5A). These results would imply feedback control on activation of *P_alpA_*, since its previously mapped transcript covers the complete region from *alpA* to *orf96323* on ICE*clc*, including *bisDC* (Fig. 1A)^25^. Although induction of BisDC was sufficient for activation of transcription from *P_alpA_*, this effectively only yielded a small subpopulation of cells with high reporter fluorescence values (Fig. 5B, C), in contrast to induction of the larger cloned gene region encompassing *parA-shi-bisDC* that activated all cells (Fig. 5B, C). The feedback loop, therefore, seemed to consist of a positive forward part that includes BisDC (reinforced by an as yet unknown other mechanism) and a modulatory repressive branch including AlpA.

**Fig 5.**
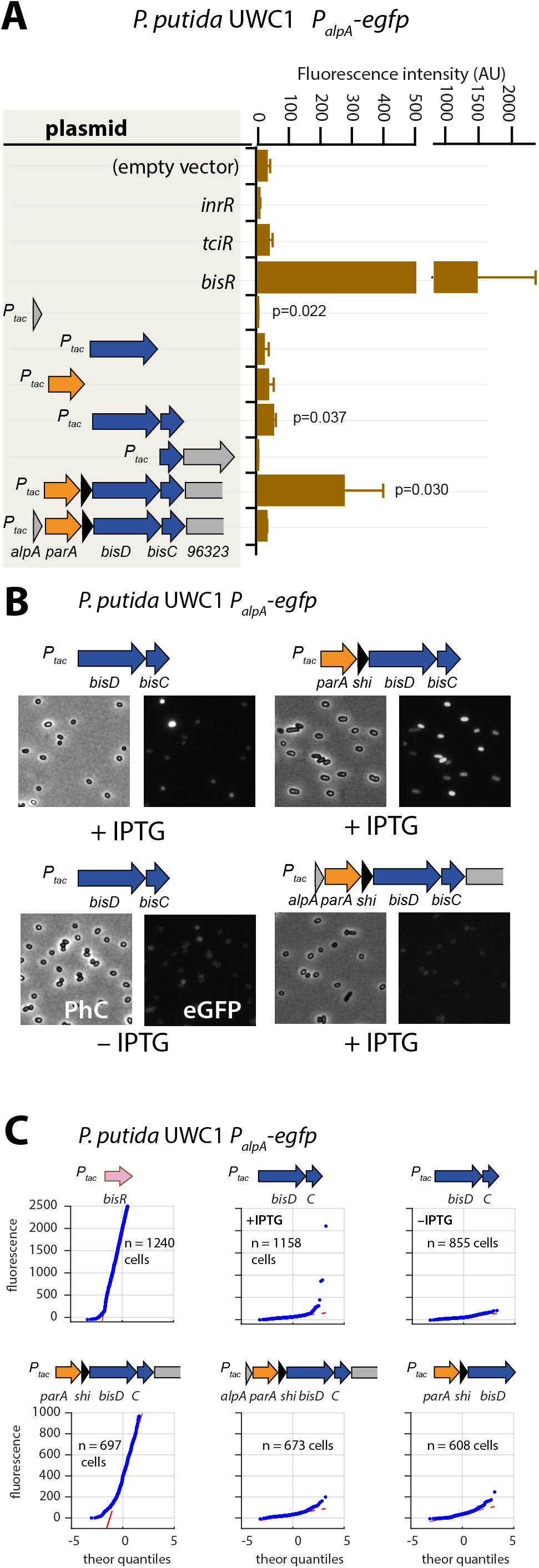
Autoregulatory feedbacks on the *alpA* promoter. **A.***P_alpA_-*reporter expression upon IPTG induction (0.05 mM) of plasmid-cloned individual genes or gene combinations (as depicted in the shaded area on the left) in *P. putida* without ICE*clc.* Bars represent means of median cell fluorescence plus one standard deviation, as in legend to Figure 2B. p-values stem from pair-wise comparisons between triplicate cultures carrying the empty vector pME6032 and the indicated plasmid-cloned gene(s). **B.** Cell images of *P. putida P_alpA_-egfp* without ICE*clc* expressing plasmid-cloned combinations with *bisDC* (fluorescence brightness scaled to 300–1200, 0.05 mM IPTG). **C.** Quantile-quantile estimation of subpopulation expression of the *P_alpA_-egfp* reporter, showing the sufficiency of *bisDC* induction for autoregulatory feedback and the reinforcement from upstream elements (*n* denotes the number of cells used for the quantile-quantile plot, summed from 6-12 images of a single replicate culture).

### Modeling suggests positive feedback loop to generate and maintain ICE*clc* bistable output

The results so far thus indicated that ICE*clc* transfer competence is initiated by TciR activating transcription of the promoter upstream of *bisR* (Fig. 6A). BisR then kickstarts expression from the *alpA-*promoter, leading to (among others) expression of BisDC. This is sufficient to induce the ‘downstream’ ICE*clc* transfer competence pathway (Fig. 1B and 6A), exemplified here by activation of the *P_int_* and *P_inR_* promoters that become exclusively expressed in the subpopulation of transfer competent cells under wild-type conditions^10^. In addition, BisDC reinforces transcription from the same *alpA-*promoter.

**Fig 6.**
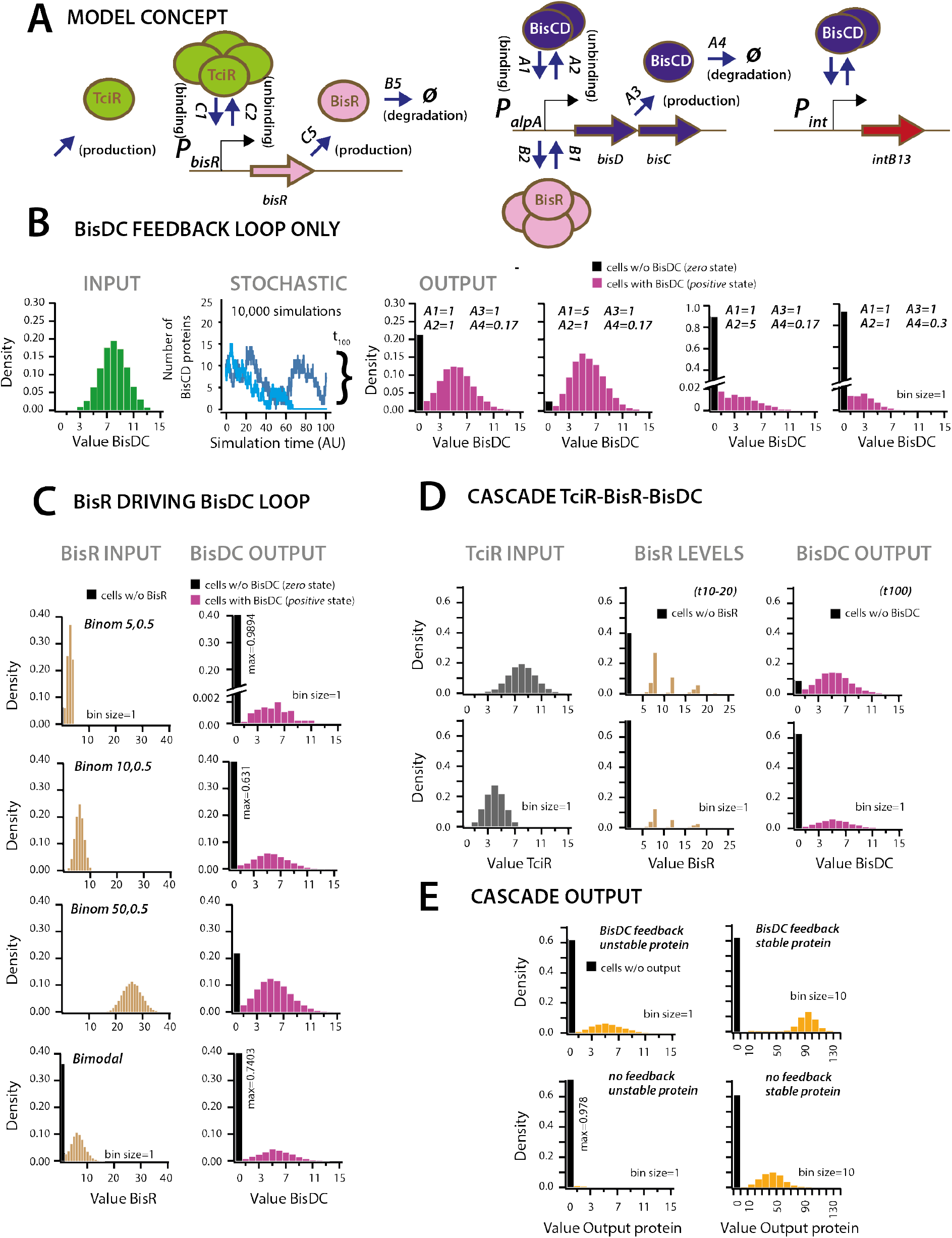
Stochastic simulations of ICE*clc* regulatory network configurations A. Conceptual model of the ICE*clc* regulatory cascade producing bistable output. Ellipses indicate the three major regulatory factors (TciR, BisR and BisDC) interacting with their target promoters (*P_bisR_*, *P_alpA_*), and BisDC-regulated downstream output (e.g., *P_int_*). Relevant simulated processes include: *production* (combination of transcription and translation, with corresponding rates: C5, A3), *oligomerization* (assumed number of protein monomers in the binding complex), *binding* and *unbinding* to the target promoter, and *degradation*. All processes are simulated as stochastic events across 100 time steps, and protein output levels are summarized from 10,000 individual stochastic simulations (detailed parametrization in Supplementary Information). **B.** Behaviour of a BisDC autoregulatory feedback loop on BisDC levels as a function of different binding (*A1*), unbinding (*A2*), production (*A3*) and degradation (*A4*) rate constants, starting from a uniformly distributed set of BisDC levels (input, in green). Black bar indicates the proportion of cells with *zero* output (i.e., non-activated circuit). **C.** As for A, but for an architecture of BisR initiating *bisDC* expression, with different input distributions (uniformly low to high, or bimodal BisR input). Note how higher or bimodal BisR input is not expected to change the median BisDC levels in active cells, but only the proportion of ‘cells’ with *positive* (magenta, bars) and *zero* state (black bars). **D.** As for A, but for the complete cascade starting with TciR. Shown are regulatory factor level distributions from two different TciR starting distributions across 10,000 simulations; for BisR integrated between time points 10 and 20, and for BisDC after 100 time steps. Bimodal expression of *zero* and *positive* states arises at the *bisR* node, but is further maintained to constant BisDC output as a result of the feedback loop. **E.** Importance of the BisDC-feedback on the output of a BisDC-dependent protein expression, dependent on its assumed protein degradation constant.

In order to understand the importance of this regulatory architecture for generating bistability, for initiating and maintaining (downstream) transfer competence, we developed a conceptual mathematical model (*Materials and methods,* SI model). The model assumes the regulatory factors TciR, BisR and BisDC, their oligomerization, as well as binding of the oligomerized forms to and unbinding from their respective nodes (i.e., the linked promoters *P_bisR_*, *P_alpA_* and *P_int_*). Binding is assumed to lead to protein synthesis and finally, protein degradation (Fig. 6A). We varied and explored the outcomes of different regulatory network architectures and parts, testing their effect on production of intermediary and downstream elements in stochastic simulations (Fig. 6A, SI model).

First we simulated the levels of BisDC in a subnetwork configuration with only BisDC activating *P_alpA_* (i.e., in absence of TciR or BisR, Fig. 6B). Stochastic simulations (n = 10,000) of this bare feedback loop with an arbitrary start of binomially distributed BisDC (mean = 8 molecules, Fig. 6B, *INPUT*), yielded a bimodal population with two BisDC output states after 100 time steps, one of which is *zero* (black bar in histograms) and the other with a mean *positive* BisDC value (magenta) (Fig. 6B). The output *zero* results when BisDC levels stochastically fall to 0 (as for the light blue line in the panel *STOCHASTIC* of Fig. 6B), since in that case there is no BisDC to stimulate its own production. Parameter variation showed that the proportion of output *zero* from the loop is dependent on the binding and unbinding constants for the *alpA* promoter, and the BisDC degradation rate (Fig. 6B, different *A1*, *A2* and *A4-*values). This simulation thus indicated that a BisDC feedback loop can produce bimodal output, once BisDC is present.

Since the feedback loop cannot start without BisDC, it is imperative to kickstart the *alpA* promoter by BisR (Fig. 6C). Simulations of a configuration that includes activation by BisR, showed how upon a single pulse of BisR, the feedback loop again leads to a bimodal population with *zero* and *positive* BisDC levels (Fig. 6C). Increasing the input level of BisR resulted in increasing the proportion of cells with *positive* BisDC state, but did not influence their mean value (Fig. 6C). Bimodally distributed BisR input also gave rise to bimodal BisCD output, but with a higher proportion of *zero* BisDC state (Fig. 6C, bimodal). In contrast to the BisDC loop alone, therefore, activation by BisR only influences the proportion of *zero* and *positive* BisDC states in the population.

In the full regulatory hierarchy of the ICE, production of BisR is controlled by TciR. Simulation of this configuration showed that bimodality already appeared at the level of BisR, and the proportions of *zero* and *positive* states of both BisR and BisDC varied depending on the relative amounts of TciR (Fig. 6D). Bimodal BisDC levels are propagated by the network architecture to downstream promoters, as a consequence of them being under BisDC control (Fig. 6E). Importantly, simulations of an architecture without the BisDC feedback loop consistently resulted in lower protein output from BisDC–regulated promoters in activated cells than with feedback (Fig. 6E). This suggests two crucial functions for the ICE regulatory network: first, to convert unimodal or stochastic (‘analog’) expression of TciR and BisR to a consistent subpopulation of cells with *positive* BisDC state, and secondly, to ensure sufficient BisDC levels to activate downstream promoters within the *positive* cell population (Fig. 6E). Through its reinforcement, bimodal expression at the *alpA*-promoter node can thus yield a stably expressed transfer competence pathway in a subpopulation of cells.

### ICE*clc* regulatory architecture exemplifies a faithful analog-to-digital converter

Simulations thus predicted that the ICE regulatory network faithfully transmits and stabilizes analog input (e.g., a single regulatory factor expressed in all cells) to bistable output (e.g., a subset of cells with transfer competence and the remainder silent). To demonstrate this experimentally, we engineered a *P. putida* without ICE*clc*, but with a single copy chromosomally inserted IPTG-inducible *bisR*, a plasmid with *parA-shi-bisDC* under control of *P_alpA_*, and a single-copy dual *P_int_-echerry* and *P_inR_-egfp* reporter (Fig. 7A). Induction from *P_tac_* by IPTG addition yields unimodal (analog) production of BisR, the mean level of which can be controlled by the IPTG concentration (Fig. S5). In the presence of all components of the system, IPTG induction of BisR led to activation of both reporters (Fig. 7B, ABC). Increasing BisR induction was converted by the feedback loop into an increased proportion of fluorescent cells (Fig. 7C). This effectively created a scalable bimodal (digital) output from unimodal input, dependent on the used IPTG concentration (Fig. 7C, Fig. S5). The proportion of fluorescent cells was in line with predictions from stochastic simulations as a function of the relative strength of *P_tac_* activation (Fig. 7D). Furthermore, in agreement with model predictions (Fig. 6C), the median fluorescence of activated cells remained the same at different IPTG (and thus BisR) concentrations (Fig. 7E). These results confirmed that the feedback loop architecture transforms an analog regulatory factor concentration (BisR) into a stabilized bimodal (digital) output.

**Fig 7.**
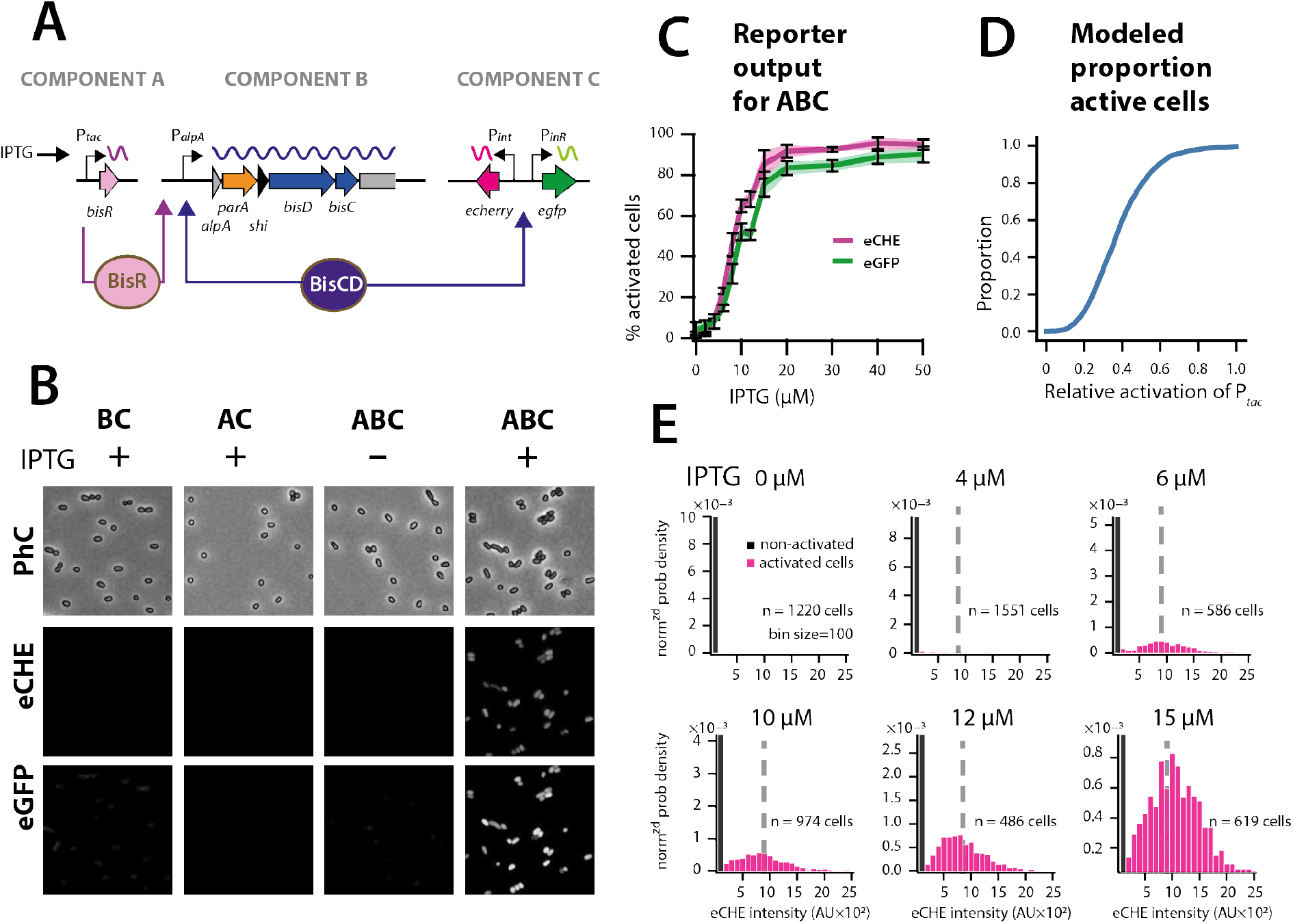
The ICE*clc* bistability generator is a scalable analog-digital converter. **A.** Schematic representation of the three ICE*clc* components used to generate scalable bistable output in *P. putida* UWC1 without ICE. **B.** Cell images of *P. putida* with the different bistability-generator components as indicated, induced in presence or absence of IPTG (0.1 mM). Fluorescence brightness scaled to between 300–1200. **C.** Proportion of active cells (estimated from quantile-quantile plotting as in Fig. 2C) as a function of IPTG concentration (same induction time for all). Lines correspond to the means from three biological replicates with transparent areas and error bars representing the standard deviation. **D.** Modeled proportion of cells with *positive* output in the architecture of Figure 6C as a function of the relative BisR starting levels from *P_tac_*. **E.** Measured distributions of eCherry fluorescence among the subpopulations of activated (magenta bars) and non-activated cells (black bars) at different IPTG concentrations, showing same subpopulation fluorescence median (dotted grey lines), as predicted in the stochastic model. AU, arbitrary units of fluorescence brightness at 500 ms exposure. *n* denotes the number of cells used to produced the histograms, summed from 6-12 images from a single replicate culture.

### BisDC-elements are widespread in other presumed ICEs

Pfam analysis detected a DUF2857-domain in the BisC protein, and further structural analysis using Phyre2 indicated significant similarities of BisC to FlhC (Fig. S1). FlhC is a subunit of the master flagellar activator FlhDC of *E. coli* and *Salmonella*^31,32^. BisD carries a ParB domain, with a predicted DNA binding domain in the C-terminal portion of the protein (Fig. S1). Although no FlhD domain was detected in BisD, in analogy to FhlDC we named the ICE*clc* activator complex BisDC, for <u>bis</u>tability regulator subunits <u>D</u> and <u>C</u>.

Database searches showed that *bisDC* loci are widespread among pathogenic and environmental *Gamma*- and *Beta-proteobacteria*, and are also found in some *Alphaproteobacteria* (Fig. S2). Phylogenetic analysis using the more distantly related sequence from *Dickeya zeae* MS2 as an outgroup indicated several clear clades, encompassing notably *bisDC* homologs within genomes of *Pseudomonas aeruginosa* and *Xanthomonas* (Fig. S6). Several genomes contained more than one *bisDC* homolog, the most extreme case being *Bordetella petrii* DSM12804 with up to four homologs belonging to four different clades (Fig. S6).

The gene synteny from *bisR* to *inrR* of ICE*clc* was maintained in several genomes (Fig. S2), suggesting them being part of related integrated ICEs. Notable exceptions included a region in *P. aeruginosa* Carb01_63, which carried an integrase gene upstream of *bisR* but that was still downstream of a *tRNA^gly^* gene (Fig. S2). This region may encompass an ICE that has retained the same integration specificity as ICE*clc* but carrying a different modular architecture where regulation and integration modules are next to each other, instead of at the opposite ends as in ICE*clc* (Fig. 1 and S2). Further exceptions included genomic regions in *P. aeruginosa* strains HS9 and W60856, which carry a gene for a LysR-type transcriptional regulator (LTTR) upstream of *bisR* with 53% and 51% amino acid identity to TciR (overlap lengths 95%), respectively. Regulation of these two elements might thus involve a *cis*-acting LTTR, rather than the *trans-*acting TciR. In the genomes of *Xanthomonas campestris* strain AW13 and *Cupriavidus nantongensis* X1 (Fig. S2), *bisD* homologs are split in two individual genes, one coding for a canonical-length ParB and the other for a BisD-homolog with only DNA-binding domain, suggesting that *bisD* on ICE*clc* may have arisen from a gene fusion.

## Discussion

ICEs operate a dual life style in their host, which controls their overall fitness as a sum of vertical descent (i.e., maintenance of the integrated state and replication with the host chromosome) and horizontal transfer (i.e., excision from the host cell, transfer and reintegration into a new host)^11,16,20^. The decision for horizontal transfer is costly and potentially damages the host cell^11,26^, which is probably why its frequency of occurrence in most ICEs is fairly low (<10^−5^ per cell in a host cell population)^20^. Consequently, the mechanisms that initiate and ensure ICE horizontal transfer must have been selected to operate under extremely low opportunity with high success. In other words, they have been selected to maximize faithful maintenance of transfer competence development, once this process has been triggered in a host cell. One would thus expect such mechanisms to impinge on rare, perhaps stochastic cellular events, yielding robust output despite cellular gene expression and pathway noise. ICE*clc* is further particular in the sense that its transfer competence is initiated in cells during stationary phase conditions^33^, which restricts global transcription and activity, and may even profoundly alter the cytoplasmic state of the cell^34^.

The results of our work here reveal that the basis for initiation and maintenance of ICE*clc* transfer competence in a minority of cells in a stationary phase population^14^, originates in a multinode regulatory network that further includes a positive feedback loop. Genetic dissection, epistasis experiments and expression of individual components in *P. putida* devoid of the ICE showed that the network consists of a number of regulatory factors, composed of MfsR, TciR, BisR and BisDC, acting sequentially on singular (TciR, BisR) or multiple nodes (BisDC). The network has an ‘upstream’ branch controlling the initiation of transfer competence, a ‘bistability generator’ that confines the input signal, and maintains the ‘downstream’ path of transfer competence to a dedicated subpopulation of cells (Fig. 1B).

The previously characterized *mfsR-marR-tciR* operon^26^, whose transcription is controlled through autorepression by MfsR, is probably the main break on activation of the upstream branch. This was concluded from effects of deleting *mfsR*, which resulted in overexpression of TciR, and massively increased and deregulated ICE transfer even in exponentially growing cells^26^. We showed here that TciR activates the transcription of a hitherto unrecognized transcription factor gene named *bisR*, but not of any further critical ICE*clc* promoters. Autorepression by MfsR in wild-type ICE*clc* results in low unimodal transcription from *P_mfsR_*^26^ and therefore, likely, to low TciR levels in all cells. TciR appeared here as a weak activator of the *bisR* promoter, suggesting that only in a small proportion of cells it manages to trigger *bisR* transcription, as our model simulations further attested.

The BisR amino acid sequence revealed only very weak homology to known functional domains, thus making it the prototype of a new family of transcriptional regulators. In contrast to TciR, BisR was a very potent activator of its target, the *alpA* promoter. Model simulations suggested that BisR triggers and transmits the response in a scalable manner to the bistability generator, encoded by the genes downstream of *P_alpA_*. Triggering of *P_alpA_* stimulated expression of (among others) two consecutive genes *bisD* and *bisC*, which code for subunits of an activator complex that weakly resembles the known regulator of flagellar synthesis FlhDC^31,32^. BisDC production was sufficient to activate the previously characterized bistable ICE*clc* promoters *P_int_* and *P_inR_*, making it the key regulator for the ‘downstream’ branch (Fig. 1B). Importantly, BisDC was also part of a feedback mechanism activating transcription from *P_alpA_*, and therefore, regulates its own production. Simulations and experimental data indicated that the feedback loop acts as a scalable analog-to-digital converter, transforming any *positive* input received from BisR into a dedicated cell that can regenerate sufficiently high BisDC levels to activate the complete downstream transfer competence pathway.

Bistable gene network architectures are characterized by the fact that expression variation is not resulting in a single mean phenotype, but can lead to two (or more) stable phenotypes - mostly resulting in individual cells displaying either one or the other phenotype^35–37^. Importantly, such bistable states are an epigenetic result of the network functioning and do not involve modifications or mutations on the DNA^38,39^. Bistable phenotypes may endure for a particular time in individual cells and their offspring, or erode over time as a result of cell division or other mechanism, after which the ground state of the network reappears. One can thus distinguish different steps in a bistable network: (i) the bistability *switch* that is at the origin of producing the different states, (ii) a *propagation* or *maintenance* mechanism and (iii) a *degradation* mechanism [11].

Some of the most well characterized bistable processes in bacteria include competence formation and sporulation in *Bacillus subtilis*^37^. Differentiation of vegetative cells into spores only takes place when nutrients become scarce or environmental conditions deteriorate^3,40^. Sporulation is controlled by a set of feedback loops and protein phosphorylations, which culminate in levels of the key regulator SpoOA~P being high enough to activate the sporulation genes^37^. In contrast, bistable competence formation in *B. subtilis* is generated by feedback transcription control from the major competence regulator ComK. Stochastic variations among ComK levels in individual cells, ComK degradation and inhibition by ComS, and noise at the *comK* promoter determine the onset of *comK* transcription, which then reinforces itself because of the feedback mechanism^41,42^. Initiation and maintenance of the ICE*clc* transfer competence pathway thus resembles DNA transformation competence in *B. subtilis* in its architecture of an auto-feedback loop (BisDC vs ComK). However, the switches leading to bistability are different, with ICE*clc* depending on a hierarchy of transcription factors (MfsR, TciR and BisR), and transformation competence being a balance of ComK degradation and inhibition of such degradation^41,42^. ICE*clc* bistability architecture is clearly different from the well-known double negative feedback control exerted by, e.g., the phage lambda lysogeny/lytic phase decision in *E. coli*^9,43^. That switch entails essentially a balance of the counteracting transcription factors CI, CII and Cro^9,43^. Interestingly, other ICEs of the SXT/R391 family carry this typical double negative feedback loop architecture, which may therefore control their (bistable) activation^23,44–46^.

Mathematically speaking, the ICE*clc* transfer competence regulatory architecture has two states, one of which is *zero* (inactive) and the other with a *positive* value (activation of transfer competence). Stochastic modeling suggested that the feedback loop maintains *positive* output during a longer time period than in its absence (although it will drop to *zero* at infinite time). Previous experimental data suggested that the tc cells indeed do not return to a silent ICE*clc* state, but become irreparably damaged, arrest their division^13^ and wither^14^. However, because their number is proportionally low, there is no fitness cost on the population carrying the ICE^11,47^. The advantage of prolonged feedback output seems that constant levels of the BisCD regulator can be maintained, allowing coordinated and organized production of the components necessary for the ICE*clc* transfer itself. This would consist of, for example, the relaxosome complex responsible for DNA processing at the origin(s) of transfer, and the mating pore formation complex^48^. Because *Pseudomonas* cells activate ICE*clc* transfer competence upon entry in stationary phase, the feedback loop may have a critical role to ensure faithful completion of the transfer competence pathway during this period of limiting nutrients, and to allow the ICE to excise and transfer from tc cells once new nutrients become available^11^.

Although our results were conclusive on the roles of the key regulatory factors (MfsR, TciR, BisR, BisDC), there may be further auxiliary and modulary factors, and environmental cues that influence the transfer competence network. For example, we previously found that deletions in the gene *inrR* drastically decreased ICE*clc* transfer capability by 45–fold and reduced reporter gene expression from *P_int_*^10^. Expression of InrR alone, however, did not show any direct activation of *P_int_*, *P_inR_* or *P_alpA_*, and InrR is thus unlikely to be a direct transcription activator protein. Our results also indicated that induction of AlpA may repress output from the *P_alpA_* promoter, and modulate the feedback loop that is initiated by BisR and maintained by BisDC. Furthermore, although induction of *bisDC* was sufficient to activate expression from *P_alpA_,* it was enhanced through an as yet uncharacterized mechanism involving its upstream regions. Previous results also highlighted the implication of the stationary phase sigma factor RpoS for *P_inR_* activation (Fig. 1B)^33^, which may be more generally important for other ICE*clc* regulatory promoters as well.

Phylogenetic analyses showed the different ICE*clc* regulatory loci (i.e., *bisR-alpA-bisDC-inrR*) to be widely conserved in Beta- and Gammaproteobacteria, with only few small variations in regulatory gene configurations. Most likely, these regions are part of ICE*clc*-like elements in these organisms, several of which have been detected previously^18^. The ICE-like elements in *Bordetella petrii* DSM12804 and at least one ICE*clc*-like element in *P. aeruginosa* JB2 were shown to excise from the chromosome, indicating that they are functional^49,50^. The ICE*clc* regulatory cascade for transfer competence thus seems widely conserved, controlling horizontal dissemination of this important class of bacterial conjugative elements.

## Materials and methods

### Strains and growth conditions

Bacterial strains and plasmid constructions used in this study are shortly described in Table 1 and with more detail in Table S1. Strains were routinely grown in lysogeny broth (LB Miller, Lab Logistics Group) at 30 °C for *P. putida* and 37 °C for *E. coli* in an orbital shaker incubator, and were preserved at –80 °C in LB broth containing 15 % (*v/v*) glycerol. Reporter assays and transfer experiments were carried out with cells grown in minimal media (MM)^51^ supplemented with 10 mM sodium succinate or 5 mM 3-chlorobenzoate (3-CBA). Antibiotics were used at the following concentrations: ampicillin (Ap), 100 μg ml^−1^ for *E. coli* and 500 μg ml^−1^ for *P. putida*; gentamycin (Gm), 10 μg ml^−1^ for *E. coli*, 20 μg ml^−1^ for *P. putida*; kanamycin (Kn), 50 μg ml^−1^; tetracycline (Tc), 12 μg ml^−1^ for *E. coli*, 100 μg ml^−1^ or 12.5 μg ml^−1^ for *P. putida* grown in LB or MM, respectively. Genes were induced from *P_tac_* by supplementing cultures with 0.05 mM isopropyl β-D-1-thiogalactopyranoside (IPTG; or else at the indicated concentrations).

**Table 1.**
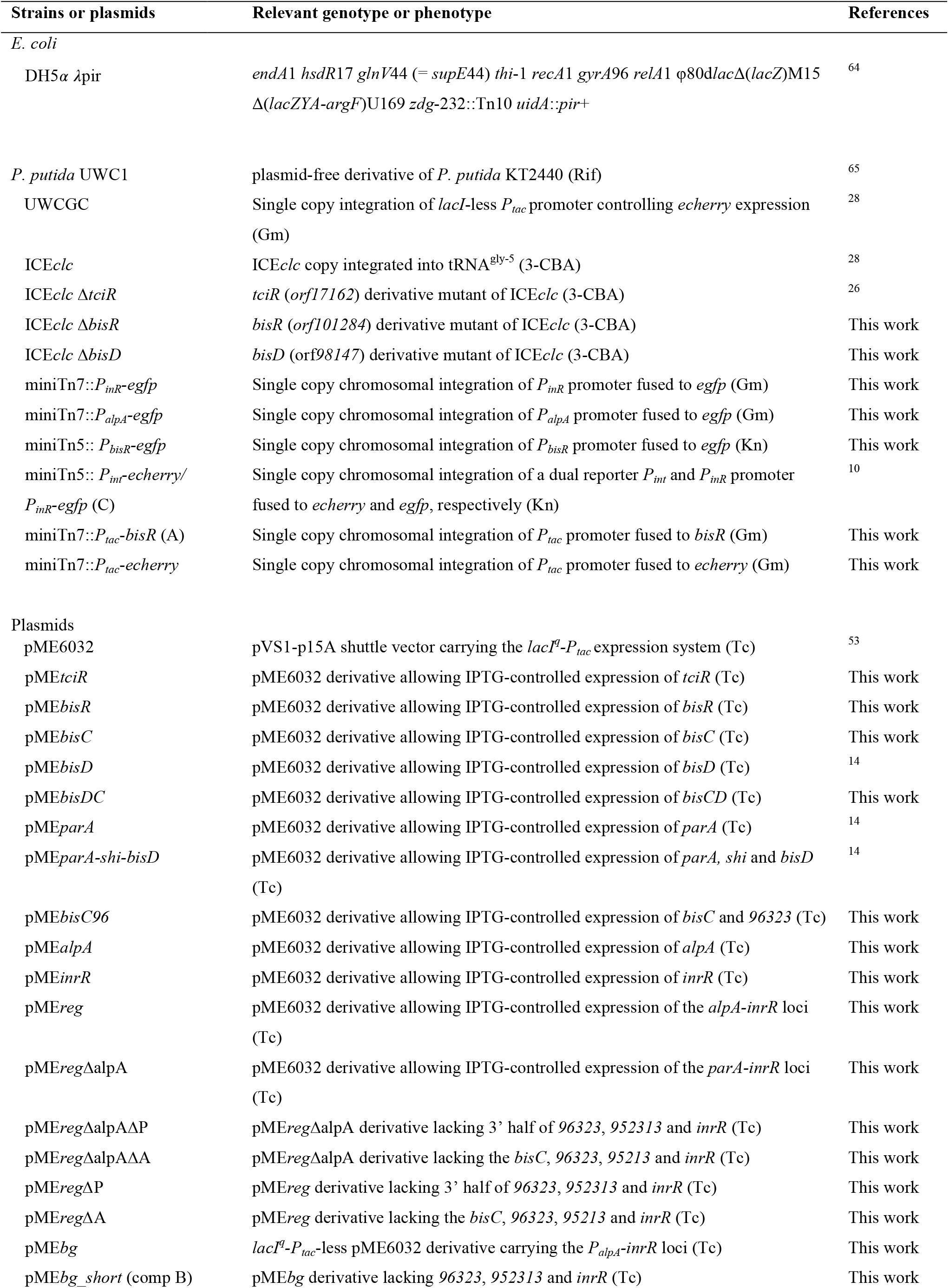
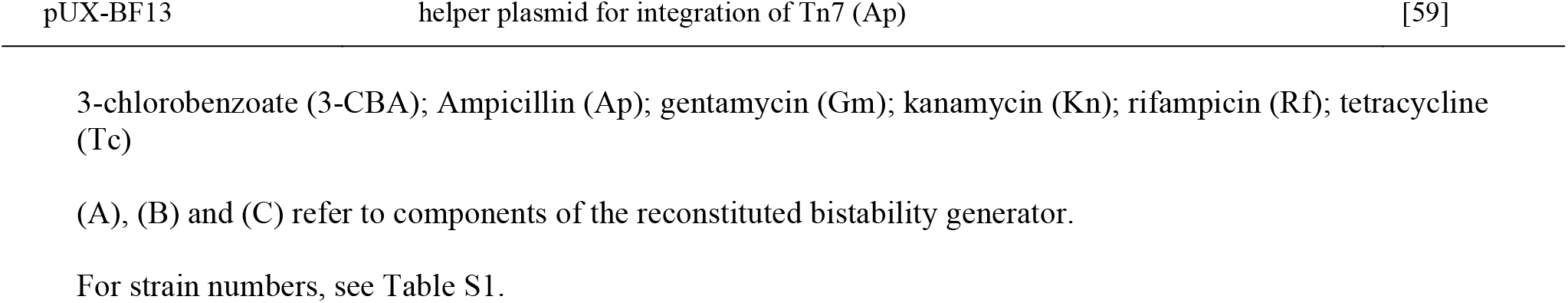
Strains and plasmids used in this study.

### Molecular biology methods

Plasmid DNA was purified using the Nucleospin Plasmid kit (Macherey-Nagel) according to manufacturer’s instructions. All enzymes used in this study were purchased from New England Biolabs. PCR reactions were carried out with primers described in Table S2. PCR products were purified using Nucleospin Gel and PCR Clean-up kits (Macherey-Nagel) according to manufacturer’s instructions. *E. coli* and *P. putida* were transformed by electroporation as described by Dower *et al*.^52^ in a Bio-Rad GenePulser Xcell apparatus set at 25 μF, 200 V and 2.5 kV for *E. coli* and 2.2 kV for *P. putida* using 2-mm gap electroporation cuvettes (Cellprojects). All constructs were verified by DNA sequencing (Eurofins).

### Plasmids and strains construction

Different ICE*clc* gene configurations were cloned in *P. putida* with or without ICE*clc*, and further with different promoter-reporter fusions, using the broad host-range vector pME6032, allowing IPTG-controlled expression from the LacI^q^-*P_tac_* promoter^53^ (Table 1). Genes *tciR, bisR, bisC, bisDC, bisC+96323*, *alpA* and *inrR* were amplified using primer pairs as specified in Table S2, with genomic DNA of *P. putida* UWC1-ICE*clc* as template. Amplicons were digested by EcoRI and cloned into EcoRI-digested pME6032 using T4 DNA ligase, producing after transformation the plasmids listed in Table 1. The 6.4-kb ICE*clc* left-end fragment encompassing *parA-inrR* was recovered from pTCB177^29^ and cloned into pME6032 (producing pME*regΔalpA*, Table S1). An *alpA-parA-shi-bisD’* fragment was amplified by PCR (Table S2) and cloned into pME6032 using EcoRI restriction sites (Table S1). The resulting plasmid was digested with SalI and the 4.8-kb fragment containing the *P_tac_* promoter, *alpA-parA-shi-bisD’* was recovered and used to replace the *parA-shi-parB* part of pME*regΔalpA*. This generated a cloned fragment encompassing *alpA* all the way to *inrR* (pME*reg*, Table S1). Further 3’ deletions removing *orf96323-inrR* or *bisC-inrR* were generated by PstI and AfeI digestion and religation (Table S1). A DNA fragment containing *P_alpA_, alpA, parA, shi* and the 5’ part of *bisD* was synthesized (ThermoFisher Scientific), and ligated by Quick-Fusion cloning (Bimake) into pME*reg*Δ*alpA* digested with PmlI and BamHI to remove the part containing *lacI^q^*, *P_tac_*, *parA*, *shi* and *bisD*. This plasmid was then digested by PstI to remove *orf96323-inrR* and religated (Table S1).

The promoter regions upstream of *bisR* or *alpA* were amplified in the PCR (Table S2) and cloned into the promoterless *egfp* reporter miniTn*5* delivery plasmid pBAM1^54^ or into a pUC18-derived miniTn*7* delivery plasmid^55^. The *P_inR_-egfp* insert was recovered from the miniTn*5*-based reporter system^10^ using HindIII and KpnI, and subsequently cloned into pUC18miniTn*7* digested by the same enzymes. The dual miniTn*7 P_inR_-egfp; P_int_-echerry* reporter has been described previously^10^. All reporter constructs were integrated in single copy into the chromosomal *attB*_Tn7_ site of *P. putida* by using pUX-BF13 for miniTn*7*, or randomly for miniTn*5*-based constructs^54,56^, in which case three independent clones were recovered, stored and analyzed.

Deletions of *bisR* or *bisD* in ICE*clc* were constructed using the two-step seamless chromosomal gene inactivation technique as described elsewhere^57^.

### ICEclc transfer assays

ICE*clc* transfer was tested with 24-h-succinate-grown donor and recipient cultures. Cells were harvested by centrifugation of 1 ml (donor) and 2 ml culture (recipient, Gm-resistant *P. putida* UWCGC) for 3 min at 1200 × *g*, washed in 1 ml of MM without carbon substrate, centrifuged again and finally resuspended in 20 μl of MM. Donor or recipient alone, and a donor-recipient mixture were deposited on 0.2–μm cellulose acetate filters (Sartorius) placed on MM agar plates, and incubated at 30 °C for 48 h. The cells were recovered from the filters in 1 ml of MM and serially diluted before plating. Donors, recipients and exconjugants were selected on MM agar plates containing appropriate antibiotics and/or carbon source (3-CBA). Transfer frequencies are reported as the mean of the exconjugant colony forming units compared to that of the donor in the same assay.

### Molecular phylogenetic analysis

BisDC phylogeny was inferred from 148 aligned homolog amino acid sequences by using the Maximum Likelihood method based on the Tamura-Nei model^58^, eliminating positions with less than 95% site coverage. The final dataset was aligned using MEGA7^59^ and contained a total of 2091 positions. Initial tree(s) for the heuristic search were obtained automatically by applying Neighbor-Joining and BioNJ algorithms to a matrix of pairwise distances estimated using the Maximum Composite Likelihood (MCL) approach, and then selecting the topology with superior log likelihood value.

### Fluorescent reporter assays

For quantification of eGFP and eCherry fluorescence in single cells, *P. putida* strains were cultured overnight at 30 °C in LB medium. The overnight culture was diluted 200 fold in 8 ml of MM supplemented with succinate (10 mM) and appropriate antibiotic(s), and grown at 30 °C and 180 rpm to stationary phase. 150 μl of culture were then sampled, vortexed for 30 seconds at max speed, after which drops of 5 μl were deposited on a regular microscope glass slide (VWR) coated with a thin film of 1% agarose in MM. Cells were covered with a 24 × 50 mm cover slip (Menzel-Gläser) and imaged immediately with a Zeiss Axioplan II microscope equipped with a 100× Plan Achromat oil objective lens (Carl Zeiss), and a SOLA SE light engine (Lumencor). A SPOT Xplorer slow-can charge coupled device camera (1.4 Megapixels monochrome w/o IR; Diagnostic Instruments) fixed on the microscope was used to capture images. Up to ten images at different positions were acquired using Visiview software (Visitron systems GMbH), with exposures set to 40 ms (phase contrast, PhC) and 500 ms (eGFP and eCherry). Cells were automatically segmented on image sets using procedures described previously^11^, from which their fluorescence (eGFP or eCherry) was quantified. Subpopulations of tc cells were quantified using quantile-quantile-plotting as described previously^27^. Fluorescent images for display were scaled to the same brightness in ImageJ^60^ as indicated, saved as 8-bit gray tiff-files and cropped to the display area in Adobe Photoshop (Adobe, 2020).

### Statistical analysis

Fluorescent reporter intensities were compared among biological triplicates. In case of mini-Tn*5* insertions, this involved three clones with potentially different insertion sites, each measured individually. For mini-Tn*7* inserted reporter constructs, we measured three biological replicates of a unique clone. Expression differences between mutants and a strain with the same genetic background but carrying the empty pME6032 plasmid were tested on triplicate means of individual median values in a one-sided t-test (the hypothesis being that the mutant expression is higher than the control). Coherent simultaneous data series were tested for significance of reporter expression or transfer frequency differences in ANOVA, followed by a post-hoc Tukey test. Quantile-quantile plots were produced in MatLab (v 2016a), violin - boxplots by using *ggplot2* in R.

### Mathematical model of ICEclc activation

ICE*clc* activation was simulated as a series of stochastic events in different network configurations (as schematically depicted in Fig. 6A, SI model). TciR, BisR, BisDC and protein output levels were then simulated using the Gillespie algorithm^61^ ^62^, implemented in *Julia* using its DifferentialEquations.jl package^63^. 10,000 individual simulations were conducted per network configuration during 100 time steps, during or after which the remaining protein levels were counted and summarized. The code for the mathematical implementation is provided in the Supplementary Information (SI code).

## Acknowledgments

The authors thank Fabrice Merz and Noëmie Matthey for their help in technical parts of this study. We thank Aleksandar Vjestica, Roxane Moritz and Andrea Daveri for critical reading. The work was supported by Swiss National Science Foundation grant to JvdM 31003A_175638 and by a SystemsX.ch Interdisciplinary grant to CM and JvdM. The funders had no role in study design, data collection and analysis, decision to publish, or preparation of the manuscript.

## Supporting Information Legends

**Fig S1. Protein domain predictions in the ICE***clc* **bistability regulators BisR, BisD and BisC.**

Genes are schematically represented by arrows with indicated size (nucleotides). Corresponding predicted protein domains using Phyre2 (prediction confidence within brackets) are shown below the respective genes, with amino acid positions indicated.

**Fig. S2 Gene organization of ICE***clc***-related regulatory loci in different bacterial species**. Diagrams show gene organization (colored arrows, to scale) of the ICE*clc* left-end regulatory loci (*Pseudomonas knackmussii* B13, Accession number HG322950.1), and homologs in *Pseudomonas aeruginosa* Carb01_63 (CP011317.1), *Herminiimonas arsenicoxydans* (CU207211.1); *Acidovorax* sp. JS42 (CP000539.1); *Burkholderia cenocepacia* ST32 (CP011917.1); *Pseudomonas aeruginosa* HS9 (CP030861.1); *Xanthomonas citri* subsp. *citri* AW13 (CP009031.1); *Pseudomonas aeruginosa* W60856 (CP008864.2); and *Cupriavidus nantongensis* X1 (CP014844.1). Genes of unknown function (dark grey) inserted between *orfs96323* and *95213* in strains AW13, HS9 (2) and W60856. Numbers below genes indicate the percentage of amino acid identity and coverage to the corresponding one from ICE*clc*. ni: no identity.

**Fig. S3. Effect of***bisDC* **containing plasmid constructs on fluorescent protein expression from a single copy dual***P_int_***-***P_inR_* **reporter in***P. putida* **without ICE***clc.*

Diagrams show distributions of individual cell fluorescence of *P. putida* from *P_int_* (eCherry) and *P_inR_* (eGFP), complemented with the indicated plasmid constructs (on the right), upon induction with 0.05 mM IPTG. Distributions represented by box plots (black on white, with first quartile, median and second quartile), underlaid with a violin distribution of all data (salmon red). Single asterisks indicate statistically significant differences of median values compared to *P. putida* carrying the empty vector in pair-wise t-test of biological triplicates (p<0.05), whereas double asterisks indicate statistically significant differences of the measured subpopulations in quantile-quantile plots.

**Fig. S4. Effect of***bisDC* **expression on reporter expression from***P_int_* **in***P. putida* **carrying ICE***clc* **with a***bisD* **deletion.**

Violin/box plots (as in legend to Fig. S3) show distributions of log10 individual cell fluorescence from *P_int_* (eCherry) of *P. putida* with ICE*clc–ΔbisD* or wild-type, complemented with the indicated plasmid constructs (in the middle), upon induction with 0.05 mM IPTG. B. Corresponding quantile-quantile plots to identify subpopulations of reporter-expressing cells. Single asterisks indicate statistically significant differences of median values compared to *P. putida* carrying the empty vector in pair-wise t-test of biological triplicates (p<0.05).

**Fig. S5. Scalable analog (***P_tac_* **) to digital bimodal (***P_int_***) expression in***P. putida* **with ICE***clc* **bistability generator.**

**A.** Unimodal eCherry expression among individual *P. putida* cells from a single copy *P_tac_*-*echerry* chromosomal insertion, as a function of increasing IPTG concentration. **B.** Bimodal eCherry fluorescence from *P_int_* among individual *P. putida* cells expressing BisR from *P_tac_* at different IPTG concentrations and carrying the plasmid cloned *P_alpA_-inrR* gene region (bistability generator). Shown are quantile-quantile plots of the observed versus expected distribution of fluorescence levels (n = number of observed cells summed from 10-12 technical replicates).

**Fig. S6. (continued from previous page) BisDC homologs among***Proteobacteria*. Maximum Likelihood phylogenetic tree of BisDC homologs among *Proteobacteria* based on amino acid alignment using the Tamura-Nei model. Numbers next to branching points indicate the percentage of trees with clustering as shown among 100 bootstraps. BisDC representatives displayed in Fig. S2 are highlighted. *C. nantogensis* X1 sequences were linked from its two separate genes.

**SI Table 1: Plasmid and used strain numbers of***P. putida* **derivatives**

**SI Table 2: Used primers in the study.**

**SI model**

ICE*clc* transfer competence bistable regulatory model REACTIONS

A: BisDC feedback loop on alpA-promoter

B: BisDC feedback loop on alpA-promoter, initiated by BisR

C: TciR activating bisR-promoter

D: BisDC activating three downstream (late) promoters

E: BisDC activating GFP reporter.

**SI code**

ICE stochastic model: code for implementation in Julia

## References

1. Shu, C. C., Chatterjee, A., Dunny, G., Hu, W. S. & Ramkrishna, D. Bistability versus bimodal distributions in gene regulatory processes from population balance. PLoS Comput Biol 7, e1002140 (2011).

2. Norman, T. M., Lord, N. D., Paulsson, J. & Losick, R. Stochastic switching of cell fate in microbes. Annu Rev Microbiol 69, 381–403 (2015).

3. Veening, J. W., Smits, W. K. & Kuipers, O. P. Bistability, epigenetics, and bet-hedging in bacteria. Annu Rev Microbiol 62, 193–210 (2008).

4. Xi, H., Duan, L. & Turcotte, M. Point-cycle bistability and stochasticity in a regulatory circuit for *Bacillus subtilis* competence. Math Biosci 244, 135–147 (2013).

5. Schultz, D., Ben Jacob, E., Onuchic, J. N. & Wolynes, P. G. Molecular level stochastic model for competence cycles in *Bacillus subtilis*. Proc Natl Acad Sci U S A 104, 17582–17587 (2007).

6. Lewis, K. Persister cells, dormancy and infectious disease. Nat Rev Microbiol 5, 48–56 (2007).

7. Chin, C. Y. et al. A high-frequency phenotypic switch links bacterial virulence and environmental survival in *Acinetobacter baumannii*. Nat Microbiol 3, 563–569 (2018).

8. Sepulveda, L. A., Xu, H., Zhang, J., Wang, M. & Golding, I. Measurement of gene regulation in individual cells reveals rapid switching between promoter states. Science 351, 1218–1222 (2016).

9. Arkin, A., Ross, J. & McAdams, H. H. Stochastic kinetic analysis of developmental pathway bifurcation in phage lambda-infected *Escherichia coli* cells. Genetics 149, 1633–1648 (1998).

10. Minoia, M. et al. Stochasticity and bistability in horizontal transfer control of a genomic island in *Pseudomonas*. Proc Natl Acad Sci U S A 105, 20792–20797 (2008).

11. Delavat, F., Mitri, S., Pelet, S. & van der Meer, J. R. Highly variable individual donor cell fates characterize robust horizontal gene transfer of an integrative and conjugative element. Proc Natl Acad Sci U S A 113, E3375–3383 (2016).

12. Delavat, F., Moritz, R. & van der Meer, J. R. Transient replication in specialized cells favors transfer of an Integrative and Conjugative Element. mBio 10(2019).

13. Takano, S., Fukuda, K., Koto, A. & Miyazaki, R. A novel system of bacterial cell division arrest implicated in horizontal transmission of an integrative and conjugative element. PLoS Genet 15, e1008445 (2019).

14. Reinhard, F., Miyazaki, R., Pradervand, N. & van der Meer, J. R. Cell differentiation to “mating bodies” induced by an integrating and conjugative element in free-living bacteria. Curr Biol 23, 255–259 (2013).

15. Waldor, M. K., Tschape, H. & Mekalanos, J. J. A new type of conjugative transposon encodes resistance to sulfamethoxazole, trimethoprim, and streptomycin in *Vibrio cholerae* O139. J Bacteriol 178, 4157–4165 (1996).

16. Johnson, C. M. & Grossman, A. D. Integrative and Conjugative Elements (ICEs): What they do and how they work. Annu Rev Genet 49, 577–601 (2015).

17. Burrus, V., Pavlovic, G., Decaris, B. & Guédon, G. Conjugative transposons: the tip of the iceberg. Mol Microbiol 46, 601–610 (2002).

18. Miyazaki, R. et al. Comparative genome analysis of *Pseudomonas knackmussii* B13, the first bacterium known to degrade chloroaromatic compounds. Environ Microbiol 17, 91–104 (2015).

19. Zamarro, M. T., Martin-Moldes, Z. & Diaz, E. The ICE*XTD* of *Azoarcus* sp. CIB, an integrative and conjugative element with aerobic and anaerobic catabolic properties. Environ Microbiol 18, 5018–5031 (2016).

20. Delavat, F., Miyazaki, R., Carraro, N., Pradervand, N. & van der Meer, J. R. The hidden life of integrative and conjugative elements. FEMS Microbiol Rev 41, 512–537 (2017).

21. Wozniak, R. A. & Waldor, M. K. Integrative and conjugative elements: mosaic mobile genetic elements enabling dynamic lateral gene flow. Nat Rev Microbiol 8, 552–563 (2010).

22. Carraro, N. & Burrus, V. Biology of three ICE families: SXT/R391, ICEBs1, and ICESt1/ICESt3. Microbiol Spectr 2, MDNA3–0008-2014 (2014).

23. Poulin-Laprade, D. & Burrus, V. A lambda Cro-like repressor is essential for the induction of conjugative transfer of SXT/R391 elements in response to DNA damage. J Bacteriol 197, 3822–3833 (2015).

24. Bellanger, X., Morel, C., Decaris, B. & Guedon, G. Derepression of excision of integrative and potentially conjugative elements from *Streptococcus thermophilus* by DNA damage response: implication of a cI-related repressor. J Bacteriol 189, 1478–1481 (2007).

25. Gaillard, M. et al. Transcriptome analysis of the mobile genome ICE*clc* in *Pseudomonas knackmussii* B13. BMC Microbiol 10, 153 (2010).

26. Pradervand, N. et al. An operon of three transcriptional regulators controls horizontal gene transfer of the Integrative and Conjugative Element ICE*clc* in *Pseudomonas knackmussii* B13. PLoS Genet 10, e1004441 (2014).

27. Reinhard, F. & van der Meer, J. R. Improved statistical analysis of low abundance phenomena in bimodal bacterial populations. PLoS ONE 8, e78288 (2013).

28. Miyazaki, R. & van der Meer, J. R. A dual functional origin of transfer in the ICE*clc* genomic island of *Pseudomonas knackmussii* B13. Mol Microbiol 79, 743–758 (2011).

29. Sentchilo, V. S., Zehnder, A. J. B. & van der Meer, J. R. Characterization of two alternative promoters and a transcription regulator for integrase expression in the *clc* catabolic genomic island of *Pseudomonas* sp. strain B13. Mol Microbiol 49, 93–104 (2003).

30. Kelley, L. A., Mezulis, S., Yates, C. M., Wass, M. N. & Sternberg, M. J. The Phyre2 web portal for protein modeling, prediction and analysis. Nat Protoc 10, 845–858 (2015).

31. Claret, L. & Hughes, C. Functions of the subunits in the FlhD(2)C(2) transcriptional master regulator of bacterial flagellum biogenesis and swarming. J Mol Biol 303, 467–478 (2000).

32. Liu, X. & Matsumura, P. The FlhD/FlhC complex, a transcriptional activator of the *Escherichia coli* flagellar class II operons. J Bacteriol 176, 7345–7351 (1994).

33. Miyazaki, R. et al. Cellular variability of RpoS expression underlies subpopulation activation of an integrative and conjugative element. PLoS Genet 8, e1002818 (2012).

34. Parry, B. R. et al. The bacterial cytoplasm has glass-like properties and is fluidized by metabolic activity. Cell 156, 183–194 (2014).

35. Ferrell, J. E., Jr. Bistability, bifurcations, and Waddington’s epigenetic landscape. Curr Biol 22, R458–466 (2012).

36. Ferrell, J. E., Jr. Self-perpetuating states in signal transduction: positive feedback, double-negative feedback and bistability. Curr Opin Cell Biol 14, 140–148 (2002).

37. Dubnau, D. & Losick, R. Bistability in bacteria. Mol Microbiol 61, 564–572 (2006).

38. Kussell, E. & Leibler, S. Phenotypic diversity, population growth, and information in fluctuating environments. Science 309, 2075–2078 (2005).

39. Balazsi, G., van Oudenaarden, A. & Collins, J. J. Cellular decision making and biological noise: from microbes to mammals. Cell 144, 910–925 (2011).

40. Veening, J. W., Smits, W. K., Hamoen, L. W. & Kuipers, O. P. Single cell analysis of gene expression patterns of competence development and initiation of sporulation in *Bacillus subtilis* grown on chemically defined media. J Appl Microbiol 101, 531–541 (2006).

41. Suel, G. M., Garcia-Ojalvo, J., Liberman, L. M. & Elowitz, M. B. An excitable gene regulatory circuit induces transient cellular differentiation. Nature 440, 545–550 (2006).

42. Maamar, H., Raj, A. & Dubnau, D. Noise in gene expression determines cell fate in *Bacillus subtilis*. Science 317, 526–529 (2007).

43. Bednarz, M., Halliday, J. A., Herman, C. & Golding, I. Revisiting bistability in the lysis/lysogeny circuit of bacteriophage lambda. PLoS ONE 9, e100876 (2014).

44. Bellanger, X., Morel, C., Decaris, B. & Guedon, G. Regulation of excision of integrative and potentially conjugative elements from *Streptococcus thermophilus*: role of the *arp1* repressor. J Mol Microbiol Biotechnol 14, 16–21 (2008).

45. Beaber, J. W. & Waldor, M. K. Identification of operators and promoters that control SXT conjugative transfer. J Bacteriol 186, 5945–5949 (2004).

46. Poulin-Laprade, D., Carraro, N. & Burrus, V. The extended regulatory networks of SXT/R391 integrative and conjugative elements and IncA/C conjugative plasmids. Front Microbiol 6, 837 (2015).

47. Gaillard, M., Pernet, N., Vogne, C., Hagenbüchle, O. & van der Meer, J. R. Host and invader impact of transfer of the *clc* genomic island into *Pseudomonas aeruginosa* PAO1. Proc Natl Acad Sci U S A 105, 7058–7063 (2008).

48. Carraro, N. & Burrus, V. The dualistic nature of integrative and conjugative elements. Mob Genet Elements 5, 98–102 (2015).

49. Obi, C. C. et al. The Integrative Conjugative Element *clc* (ICE*clc*) of *Pseudomonas aeruginosa* JB2. Front Microbiol 9, 1532 (2018).

50. Lechner, M. et al. Genomic island excisions in *Bordetella petrii*. BMC Microbiol 9, 141 (2009).

51. Gerhardt, P. et al. Manual of methods for general bacteriology. (American Society for Microbiology, Washington, D.C., 1981).

52. Dower, W. J., Miller, J. F. & Ragsdale, C. W. High efficiency transformation of *E. coli* by high voltage electroporation. Nucleic Acids Res 16, 6127–6145 (1988).

53. Heeb, S. et al. Small, stable shuttle vectors based on the minimal pVS1 replicon for use in gram-negative, plant-associated bacteria Mol Plant Microbe Interact 13, 232–237 (2000).

54. Martinez-Garcia, E., Calles, B., Arevalo-Rodriguez, M. & de Lorenzo, V. pBAM1: an all-synthetic genetic tool for analysis and construction of complex bacterial phenotypes. BMC Microbiol 11, 38 (2011).

55. Choi, K. H. et al. A Tn*7*-based broad-range bacterial cloning and expression system. Nat Methods 2, 443–448 (2005).

56. Koch, B., Jensen, L. E. & Nybroe, O. A panel of Tn*7*-based vectors for insertion of the *gfp* marker gene or for delivery of cloned DNA into Gram-negative bacteria at a neutral chromosomal site. J Microbiol Meth 45, 187–195 (2001).

57. Martinez-Garcia, E. & de Lorenzo, V. Engineering multiple genomic deletions in Gram-negative bacteria: analysis of the multi-resistant antibiotic profile of *Pseudomonas putida* KT2440. Environ Microbiol 13, 2702–2716 (2011).

58. Tamura, K. & Nei, M. Estimation of the number of nucleotide substitutions in the control region of mitochondrial DNA in humans and chimpanzees. Mol Biol Evol 10, 512–526 (1993).

59. Kumar, S., Stecher, G. & Tamura, K. MEGA7: Molecular Evolutionary Genetics Analysis version 7.0 for bigger datasets. Mol Biol Evol 33, 1870–1874 (2016).

60. Schneider, C. A., Rasband, W. S. & Eliceiri, K. W. NIH Image to ImageJ: 25 years of image analysis. Nat Methods 9, 671–675 (2012).

61. Gillespie, D. T. Exact stochastic simulation of coupled chemical reactions. J Phys Chem 81, 2340–2361 (1977).

62. Gillespie, D. T. A general method for numerically simulating the stochastic time evolution of coupled chemical reactions. . J Comput Phys 22, 403–434 (1976).

63. Rackauckas, C. & Nie, Q. DifferentialEquations.jl - A performant and feature-rich ecosystem for solving differential equations in Julia. J Open Res Softw 5, 15 (2017).

64. Platt, R., Drescher, C., Park, S. K. & Phillips, G. J. Genetic system for reversible integration of DNA constructs and *lacZ* gene fusions into the *Escherichia coli* chromosome. Plasmid 43, 12–23 (2000).

65. McClure, N. C., Weightman, A. J. & Fry, J. C. Survival of *Pseudomonas putida* UWC1 containing cloned catabolic genes in a model activated-sludge unit. Appl Environ Microbiol 55, 2627–2634 (1989).

